## Supplemental File for "DIETARY RESCUE OF ADULT BEHAVIORAL DEFICITS IN THE *FMR1* KNOCKOUT MOUSE"

### *Methods*

#### *Body Weight Analyses*

During the post-weaning supplementation phase, subjects receiving the two experimental dietary conditions were monitored for changes in body weight from PD21 (weaning date) through behavioral testing, until sacrifice at the conclusion of all testing. For the prenatal paradigm, body weight was assessed weekly beginning at the first week of testing (PD60) until just prior to sacrifice. For the post-weaning paradigm, the final sample sizes were as follows: Standard WT = 11, Standard KO = 10, Control Fat WT = 16, Control Fat KO = 16, Omega-3 WT = 19, Omega-3 KO = 16. The final sample size for the prenatal paradigm was as follows: Standard WT = 23, Standard KO = 12, Control Fat WT = 16, Control Fat KO = 16, Omega-3 WT = 13, Omega-3 KO = 12.

#### *Open Field*

The open field test was conducted to evaluate the effects of the three diet conditions on activity levels. Prior to testing, animals were allowed to habituate to the testing room for at least 30 minutes. The testing arena consisted of a clear acrylic box (40x40x30cm). Background noise levels in the room were then limited to 60 dB. The lighting inside the test chamber was approximately 100 lux and uniform through the chamber. Subjects were placed into the testing arena and allowed to explore freely for 30 minutes and the experimenter was not present during the testing window. During the task, activity level variables (i.e. grooming, rearing, clockwise and counterclockwise revolutions) were measured and compiled by a computer operated optical animal activity system (Fusion by AccuScan Instruments, Inc.; USA). This system also measured other exploratory behaviors such as grooming, rearing, clockwise, and counterclockwise rotations, as well as stereotypic behavior, which accounts for repeated breaking of the same set of beams, (i.e. during grooming behavior). Following testing, subjects were returned to an alternate cage until all mice in the home cage were tested, before being returned to the home cage. Between subjects, the arena was thoroughly cleaned using a 30% isopropyl alcohol solution and dried thoroughly. For the post-weaning paradigm, the final sample sizes were as follows: Standard WT = 11, Standard KO = 10, Control Fat WT = 16, Control Fat KO = 16, Omega-3 WT = 19, Omega-3 KO = 16. The final sample size for the prenatal paradigm was as follows: Standard WT = 21, Standard KO = 12, Control Fat WT = 16, Control Fat KO = 16, Omega-3 WT = 14, Omega-3 KO = 14.

*Elevated Plus Maze*

In addition to how the data was extracted from the elevated plus maze task as described in the main body of the study, these videos were also scored later offline for head dip behavior and rearing activities by an experimenter blind to the experimental condition of the subject. Following testing, the test mice were returned to an alternate cage until all mice in the home cage were tested, before being returned to the home cage. Sample sizes for this assessment were specified in main body of the study.

*Nose Poke Assay*

Nose poke behavior in a hole board arena was used as a measure of repetitive behavior. Subjects were given a 30-minute habituation period prior to testing. The testing apparatus consisted of a board with 16 equidistant holes that were 1” in diameter and approximately 0.75” in depth inserted into a clear plastic arena (40 x 40 x 30 cm). Behavior was considered a nose-poke when the subject inserts the nose as far in as the eye. During the 10-minute testing window, the number and location of these pokes was recorded by a researcher blind to the experimental condition of the subject. Following the conclusion of testing, the mice were returned to an alternate cage with other tested mice. The arena was cleaned thoroughly with 30% isopropyl alcohol between subjects. For the post-weaning paradigm, the final sample sizes were as follows: Standard WT = 10, Standard KO = 10, Control Fat WT = 16, Control Fat KO = 16, Omega-3 WT = 18, Omega-3 KO = 16. The final sample size for the prenatal paradigm was as follows: Standard WT = 24, Standard KO = 11, Control Fat WT = 16, Control Fat KO = 15, Omega-3 WT = 14, Omega-3 KO = 14.

*Social Partition*

To evaluate differences in sociability, animals were tested in the social partition paradigm. The animal was first housed overnight (approximately 24 hours) in a cage divided into two chambers by a clear partition with 0.6 cm diameter holes placed randomly, and a sex/weight/age-matched conspecific was placed in the other side. The next day the testing paradigm consisted of three testing phases. In each testing phase, the duration and frequency of visits to the partition were measured for 5 minutes. The first phase (familiar) measured the interaction at the partition with the mouse it has been previously housed with for 24 hours. In the second phase (unfamiliar), the conspecific was replaced with a novel conspecific. The last phase (familiar 2), the previous conspecific was placed behind the partition and then the behavior of the experimental subject was measured on the same constructs as the other trials. Duration and frequency was recorded by a live observer using the Ethom computer software to measure the time and frequency of the social interaction behavior (Taiwanica, 2000). For the post-weaning paradigm, the final sample sizes were as follows: Standard WT = 11, Standard KO = 10, Control Fat WT = 16, Control Fat KO = 16, Omega-3 WT = 19, Omega-3 KO = 16. The final sample size for the prenatal paradigm was as follows: Standard WT = 24, Standard KO = 12, Control Fat WT = 16, Control Fat KO = 15, Omega-3 WT = 14, Omega-3 KO = 14.

### *Results*

#### *Body Weight Analyses*

*Post-weaning paradigm.* During the post-weaning period (3 weeks of age or PD21) to 12 weeks of age (prior to behavioral testing), body weight measurements during this period were analyzed using a two-factor repeated measures ANOVA (Genotype [wildtype, *Fmr1* KO] x Diet [control fat, omega-3]) for all available data points (Figure S1A). Results indicated that *Fmr1* KOs gained weight similarly to their wildtype counterparts, $F_{genotype}(1, 57) = 0.05$, $p = 0.83$, $F_{genotype\ x\ time}(9, 513) = 0.27$, $p = 0.98$. However, dietary supplementation with omega-3 fatty acids did significantly increase body weight over time, $F_{diet}(1, 57) = 9.8$, $p = 0.003$, $F_{diet\ x\ time}(9, 513) = 2.89$, $p = 0.002$. Diet did not interact with genotype, $F_{diet\ x\ genotype}(2, 57) = 0.11$, $p = 0.74$, $F_{genotype\ x\ diet\ x\ time}(9, 513) = 0.74$, $p = 0.68$. At the final time point, final body weight was assessed prior to sacrifice for all three dietary manipulation groups for all available data points. Results indicated that post-weaning supplementation results in significantly increased body weight in animals treated with omega-3 fatty acids, compared to only the control fat condition, $F_{diet}(2, 81) = 3.20$, $p = 0.05$ (Figure S1B). Loss of *Fmr1* did not impact body weight at this time, $F_{genotype}(1, 81) = 1.49$, $p = 0.23$, $F_{diet\ x\ genotype}(2, 81) = 0.71$, $p = 0.50$.

*Prenatal paradigm.* For the prenatal paradigm, body weight was assessed beginning at the first day of testing (PD60) (Figure S1C). Results indicated no lasting effect of diet on growth prior to that time, $F_{diet}(2, 91) = 0.13$, $p = 0.88$, and diet did not significantly interact with genotype, $F_{diet\ x\ genotype}(2, 91) = 0.41$, $p = 0.67$. Loss of *Fmr1* had no effect on body weight measured at this time, $F_{genotype}(1, 91) = 0.01$, $p = 0.91$. This same finding persisted at the final time point prior to sacrifice as well (Figure S1D): $F_{genotype}(1, 91) = 0.04$, $p = 0.85$; $F_{diet}(2, 91) = 1.27$, $p = 0.29$; $F_{diet\ x\ genotype}(2, 91) = 0.70$, $p = 0.50$.

*Open Field*

*Post-weaning paradigm.* For the post-weaning paradigm, in the open field test, *Fmr1* KO mice demonstrated hyperactivity for movement time, $F_{genotype}(1, 87) = 4.63$, $p = 0.03$ (Figure S2A). However, they did not show hyperactivity on any other variable measured: total distance, $F_{genotype}(1, 87) = 0.04$, $p = 0.84$ (Figure S2B), number of rearings, $F_{genotype}(1, 87) = 3.34$, $p = 0.07$ (Figure S2C), or stereotypy time, $F_{genotype}(1, 87) = 3.66$, $p = 0.06$ (Figure S2D). These parameters were also not influenced by diet: total distance, $F_{diet}(2, 87) = 1.50$, $p = 0.23$, movement time, $F_{diet}(2, 87) = 0.82$, $p = 0.45$, number of rearings, $F_{diet}(2, 87) = 0.22$, $p = 0.80$, or stereotypy time, $F_{diet}(2, 87) = 0.09$, $p = 0.92$. Genotype and diet did not interact on any of the variables either: total distance, $F_{diet\ x\ genotype}(2, 87) = 0.61$, $p = 0.54$, movement time, $F_{diet\ x\ genotype}(2, 87) = 1.34$, $p = 0.27$, number of rearings, $F_{diet \times genotype}(2, 87) = 1.08$, $p = 0.34$, or stereotypy time, $F_{diet \times genotype}(2, 87) = 2.08$, $p = 0.13$.

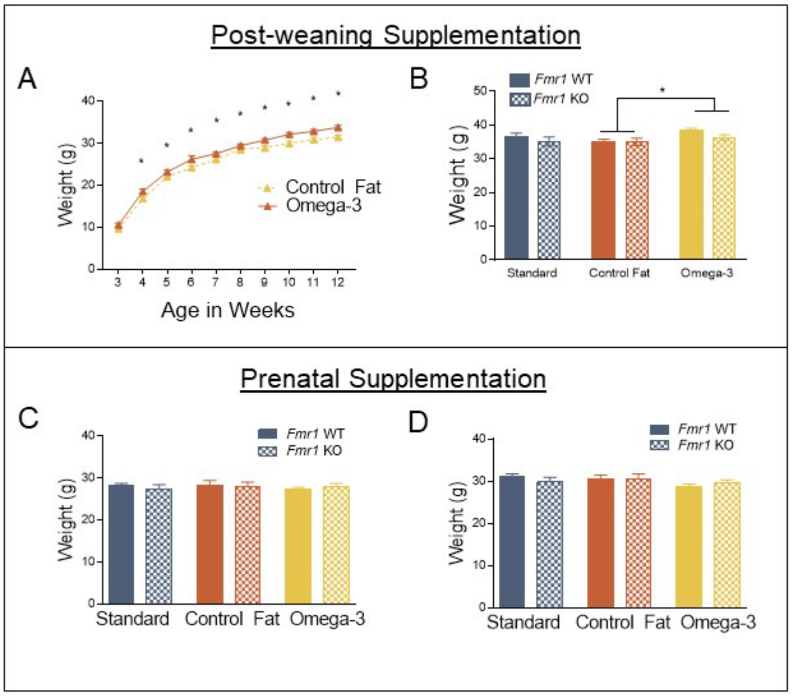


*Figure S1*. Post-weaning supplementation with omega-3 fatty acids increases body weight, while prenatal supplementation has no impact on body weight. A. Weight gain was significantly higher in the omega-3 fatty acid condition. B. Final body weight prior to sacrifice was higher in subjects receiving the omega-3 fatty acid diet. C. No lasting effect of prenatal supplementation was detected prior to behavioral testing on PD60. D. No changes in body weight were detected prior to sacrifice. Data are expressed as mean ± SEM. * = $P < 0.05$. A designation of "b" indicates that this group differed from the "a" comparison group at the level of $p < 0.05$.

*Prenatal paradigm.* For the prenatal paradigm, in the open field test, loss of *Fmr1* did not significantly impact the time spent moving during the task, $F_{genotype}(1, 87) = 3.66$, $p = 0.06$ (Figure S2E). Dietary supplementation did not impact movement time, $F_{diet}(2, 87) = 0.08$, $p = 0.93$, nor was there an interaction of genotype and diet, $F_{genotype \times diet}(2,

87) = 0.25, p = 0.78. However, some measures suggested that *Fmr1* KO mice were significantly hyperactive, showing increased total distance across the 30 minute testing period, $F_{genotype}(1, 87) = 27.35$, $p = 0.001$ (Figure S2F). Dietary supplementation did not impact distance moved, $F_{diet}(2, 87) = 0.31$, $p = 0.74$, nor was there an interaction ,$F_{genotype\ x\ diet}(2, 87) = 0.46$, $p = 0.63$. Number of rearings was not impacted by loss of *Fmr1*, $F_{genotype}(1, 87) = 1.63$, $p = 0.21$ (Figure S2G). Dietary supplementation did not impact number of rearings, $F_{diet}(2, 87) = 2.07$, $p = 0.13$, nor was there a significant interaction, $F_{genotype\ x\ diet}(2, 87) = 0.97$, $p = 0.39$. Time spent engaged in stereotypic behaviors was also not impacted by loss of *Fmr1*, $F_{genotype}(1, 87) = 0.20$, $p = 0.89$ (Figure S2H). Dietary supplementation did not impact stereotypy time, $F_{diet}(2, 87) = 0.08$, $p = 0.93$, nor was there a significant interaction, $F_{genotype\ x\ diet}(2, 87) = 0.39$, $p = 0.68$.

*Elevated Plus Maze*

*Post-weaning paradigm.* For the post-weaning paradigm, results for aspects of exploratory behavior indicated that similar to the effects on general locomotion, exposure to post-weaning omega-3 fatty acids, and to a lesser extent the control fat diet, significantly reduced exploration of the elevated plus maze. Specifically, results indicated a main effect of diet for time spent looking over the edge of the maze, $F_{diet}(2, 76) = 8.45$, $p < 0.001$, as well as the duration of rearing behavior in the closed arm of the maze, $F_{diet}(2, 76) = 7.01$, $p = 0.002$. Post-hoc analyses with LSD indicated that both the control

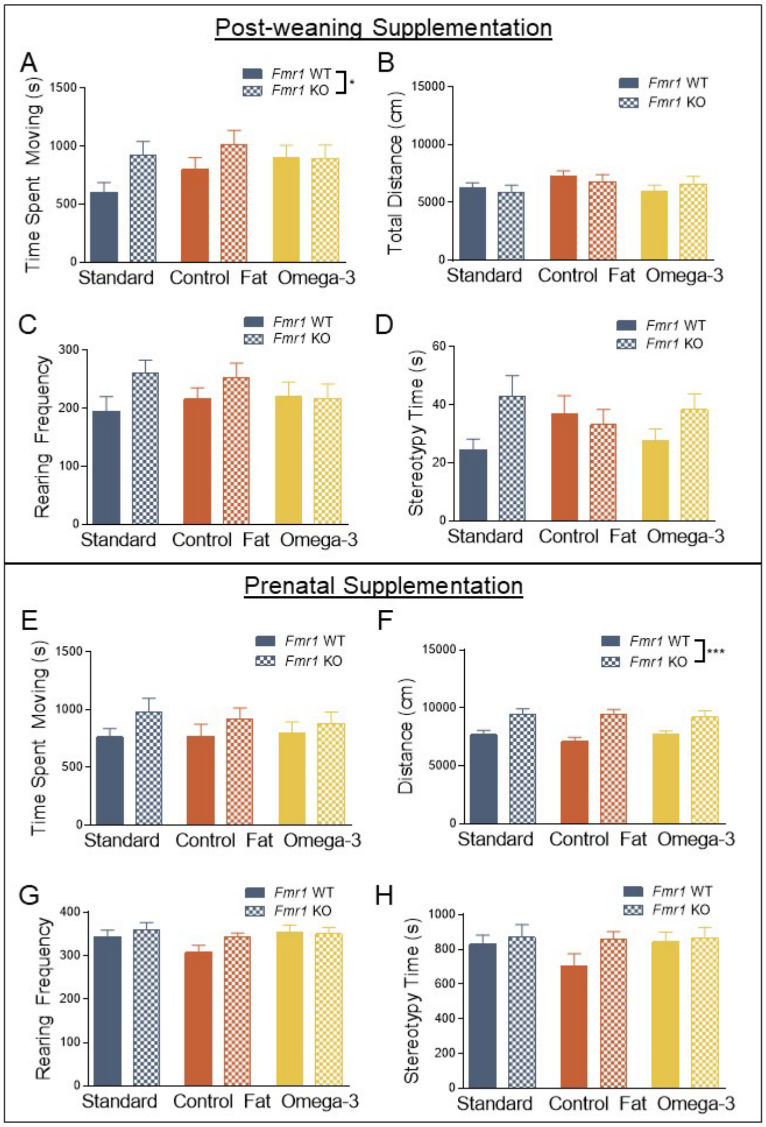


*Figure S2*. In the open field, neither prenatal nor post-weaning dietary manipulation had an impact on hyperactivity in the *Fmr1* knockout. A. *Fmr1* knockouts demonstrate hyperactivity when measuring movement time in the open field. B. Total distance moved showed no significant differences. C. Number of rearings showed no significant differences. D. Stereotypy time showed no significant differences. E. No effect was detected for movement time in the prenatal paradigm. F. *Fmr1* knockouts demonstrate hyperactivity when measuring distance moved in the open field. G. No differences were detected for rearing frequency. H. No differences were detected for stereotypy time. Data are expressed as mean ± SEM. * = P < 0.05, *** = P < 0.001. were significantly reduced compared to the standard diet.

fat ("b") and omega-3 fatty acids ("b") significantly reduced duration of head dips compared to the standard diet ("a"), at the level of $p < 0.01$ (Figure S3A). Post-hoc analyses with LSD indicated a similar pattern for closed rearing duration for both control fat ("b") and omega-3 dietary conditions ("b"), at the level of $p < 0.05$ (Figure S3B). No effect of genotype was detected for either head dip duration, $F_{genotype}(1, 76) = 0.28$, $p = 0.60$, or duration of rearings, $F_{genotype}(1, 76) = 0.39$, $p = 0.54$. Moreover, no significant interaction was detected for head dip duration, $F_{genotype \times diet}(1, 76) = 0.23$, $p = 0.80$, as well as closed rear duration, $F_{genotype \times diet}(1, 76) = 1.75$, $p = 0.18$.

*Prenatal paradigm.* For the prenatal paradigm, results for exploratory behavior in the elevated plus maze indicated that *Fmr1* KOs spent more time exploring over the sides of the open arm, $F_{genotype}(1, 84) = 4.07$, $p = 0.05$ (Figure S3C), and less time rearing in the closed arm, $F_{genotype}(1, 84) = 11.59$, $p = 0.001$ (Figure S3D). These effects were significantly attenuated by exposure to the dietary manipulations, for both head dip duration, $F_{diet}(2, 84) = 4.44$, $p = 0.02$, as well as closed rearing duration, $F_{diet}(2, 84) = 4.94$, $p = 0.01$. Post-hoc analyses with LSD indicated that this effect was only true for the omega-3 diet compared to the standard diet, $p = 0.01$. However, for the closed rear duration, both control fat ($p = 0.001$) and omega-3 ($p = 0.002$) This impact of the diet was not additive for both head dip duration, $F_{genotype \times diet}(1, 84) = 0.60$, $p = 0.55$, as well as closed rear duration, $F_{genotype \times diet}(1, 84) = 0.26$, $p = 0.77$, suggesting it did not exacerbate the phenotype. Post-hoc analyses confirmed this assertation, demonstrating that all groups differed from the Standard WT condition, but did not differ among themselves, at the level of p < 0.05.

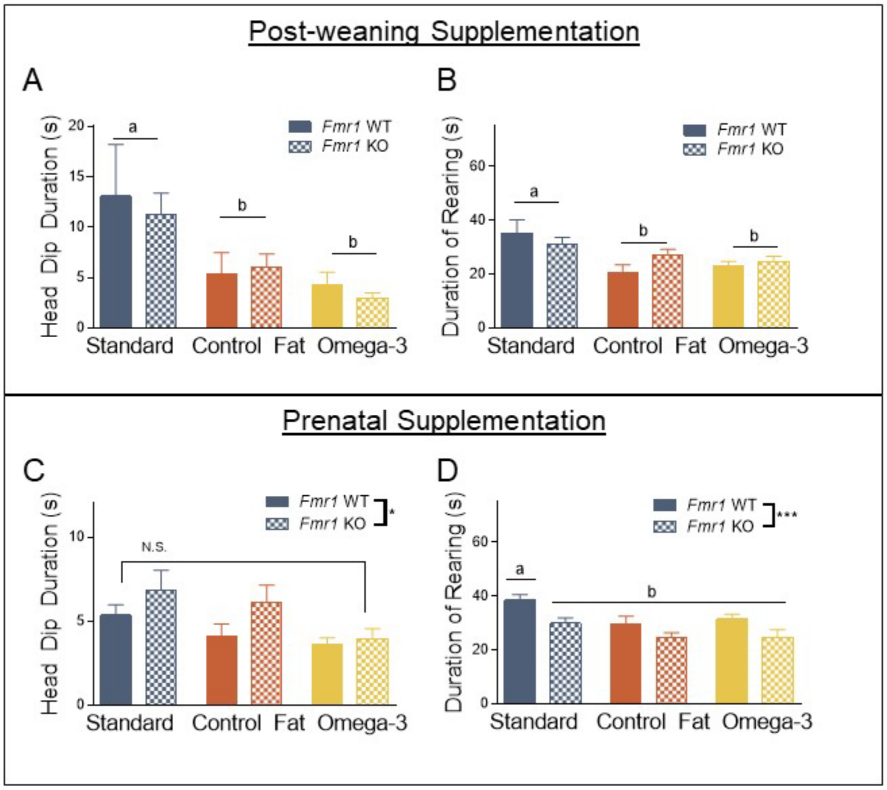


*Figure S3.* Both post-weaning and prenatal exposures to omega-3 fatty acids and the control fat diet significantly reduce exploratory behavior in the elevated plus maze. A. Post-weaning exposure to both experimental diets reduced the duration spent looking over the elevated plus maze. B. This same exposure also reduced the number of rearings in the closed arm of the maze. C. *Fmr1* knockouts spent more time looking over the side of the maze, and this was attenuated by exposure to the omega-3 fatty acid diet. D. Loss of *Fmr1* and exposure to either of the experimental diets prenatally reduced rearing behavior, however these effects were not additive. Data are expressed as mean ± SEM. * = P < 0.05, *** = P < 0.001. A designation of “b” indicates that this group differed from the “a” comparison group at the level of p < 0.05.

*Nose-Poke Assay*

*Post-weaning paradigm.* For the post-weaning paradigm, results for the latency to hole poke indicated that deletion of *Fmr1* did not reduce the mean latency, $F_{genotype}(1, 80) = 1.50$, $p = 0.22$ (Figure S4A). Diet did not impact this variable, $F_{diet}(2, 80) = 0.12$, $p = 0.89$. The combination of genotype and diet also did not impact this variable, $F_{genotype\ x\ diet}(2, 80) = 0.04$, $p = 0.97$. For total holes poked (Figure S4B), deletion of *Fmr1* did not impact this variable, $F_{genotype}(1, 80) = 1.10$, $p = 0.30$. Diet also did not significantly affect the number of holes poked overall, $F_{diet}(2, 80) = 1.86$, $p = 0.16$. Moreover, the combination of genotype and diet did not impact the number of holes poked overall, $F_{genotype\ x\ diet}(2, 80) = 0.34$, $p = 0.72$.

*Prenatal paradigm.* For prenatal paradigm, results for the latency to hole poke indicated that deletion of *Fmr1* had no impact on latency to engage in hole-poking behavior, $F_{genotype}(1, 89) = 1.03$, $p = 0.31$. However, diet significantly impacted this latency, $F_{diet}(2, 89) = 4.97$, $p = 0.01$. Further post-hoc multiple comparisons testing indicated that both control fat and omega-3 fatty acids reduced the latency to hole poke, at the level of $p < 0.05$ (Figure S4C), and magnitude of this effect was similar between the two experimental groups. The unique combination of diet and genotype did not significantly influence the latency to engage in hole-poking behavior, $F_{genotype\ x\ diet}(2, 89) = 0.38$, $p = 0.68$. Loss of *Fmr1* also did not influence the frequency of hole-poking behavior overall, $F_{genotype}(1, 89) = 2.05$, $p = 0.16$. Similar to latency, diet significantly impacted the number of hole-pokes, $F_{diet}(2, 89) = 7.71$, $p = 0.001$. Subsequent post-hoc multiple comparisons with LSD indicated that both omega-3 and control fat diets significantly increased hole-poking behavior (Figure S4D), and the magnitude of this difference was similar for both groups, at the level of $p < 0.05$. To constitute an ASD-like increase in repetitive behaviors, and not more general increases in directed exploration, we next examined whether animals indicated a preference for any particular hole or type of hole according to a previous analysis paradigm (Moy et al., 2008). Visual inspection of the data indicated that no group demonstrated “high” (> 12.5%) preference for any type of hole or specific hole (Figure S4E).

*Social Partition*

*Post-weaning paradigm.* For the post-weaning paradigm, both duration, $F_{genotype}(1, 82) = 1.02$, $p = 0.32$ (Figure S5A), and frequency, $F_{genotype}(1, 82) = 0.98$, $p = 0.32$ (Figure S5B), were unaffected by loss of *Fmr1* across all trials. When examining across the three different types of trials, loss of *Fmr1* did not interact with trial for both duration of visits, $F_{genotype\ x\ trial}(2, 164) = 1.65$, $p = 0.20$, and the number of visits to the partition, $F_{genotype\ x\ trial}(2, 164) = 0.32$, $p = 0.73$. Diet did, however, significantly impact the duration, $F_{diet}(2, 82) = 26.95$, $p = 0.0001$ (Figure S5A), and number of visits to the partition, $F_{diet}(2, 82) = 18.67$, $p = 0.0001$ (Figure S5B), across all trials. Post-hoc results for multiple comparisons indicated that both omega-3 (“b”) and control fat diets (“b”) significantly increased both the duration and frequency of visits to the partition compared to the standard diet (“a”), at the level of $p < 0.05$. This effect was consistent across all three trials, as diet did not significantly interact with trial for both duration, $F_{diet\ x\ trial}(4, 164) = 2.12$ $p = 0.08$, as well as frequency of visits to the partition, $F_{diet\ x\ trial}(4, 164) = 0.58$, $p = 0.68$. Finally, the unique combination of genotype and diet did not impact either duration, $F_{genotype\ x\ diet}(2, 82) = 0.90$, $p = 0.41$, or frequency of visits,

$F_{genotype \times diet}(2, 82) = 2.31$, $p = 0.11$, across all trials. This combination also failed to interact significantly with trial, for both duration, $F_{genotype \times diet \times time}(4, 164) = 0.40$, $p = 0.81$, as well as frequency, $F_{genotype \times diet \times time}(4, 164) = 0.54$, $p = 0.71$.

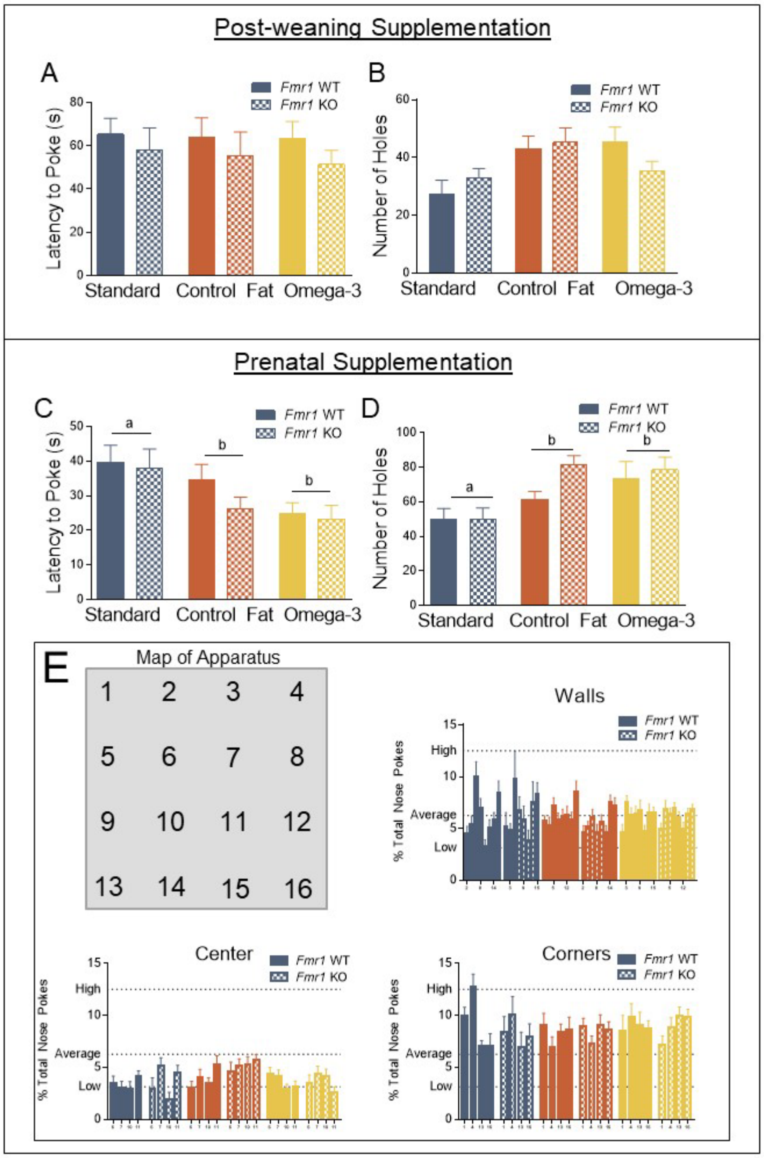


*Figure S4.* Post-weaning supplementation with high fat diets does not affect nose poking behavior, while prenatal supplementation increases directed exploration but not ASD-like repetitive behavior. A. In the post-weaning paradigm, no effects were detected for latency to the first hole-poke. B. Similarly, no effects were detected for overall number of holes poked. C. Prenatal exposure to both experimental diets reduced the latency to hole-poke. D. Similarly, prenatal exposure to the experimental diets increased the number of overall hole-pokes. E. However, when assessing for potential indicators of repetitive hole-poking, visual inspection of the data indicates no preference for a type of hole or single location. Data are expressed as mean ± SEM.

*Prenatal paradigm.* For the prenatal paradigm, both duration, $F_{genotype}(1, 89) = 0.001$, $p = 0.98$ (Figure S5C), and frequency, $F_{genotype}(1, 89) = 0.76$, $p = 0.39$ (Figure S5D), were unaffected by loss of *Fmr1* across all trials. When examining across the three different types of trials, loss of *Fmr1* did not interact with trial for both duration of visits, $F_{genotype\ x\ trial}(2, 178) = 1.04$, $p = 0.36$, and the number of visits to the partition, $F_{genotype\ x\ trial}(2, 178) = 0.09$, $p = 0.92$. Diet also did not significantly impact the duration, $F_{diet}(2, 89) = 1.44$, $p = 0.24$, or number of visits to the partition, $F_{diet}(2, 89) = 0.75$, $p = 0.48$, across all trials. Similar to genotype, diet did not significantly interact with trial for both duration, $F_{diet\ x\ trial}(4, 178) = 0.95$, $p = 0.44$, as well as frequency of visits to the partition, $F_{diet\ x\ trial}(4, 178) = 0.53$, $p = 0.71$. Finally, the unique combination of genotype and diet did not impact either duration, $F_{genotype\ x\ diet}(2, 89) = 0.38$, $p = 0.69$, or frequency of visits, $F_{genotype\ x\ diet}(2, 89) = 0.02$, $p = 0.98$, across all trials. This combination also failed to interact significantly with trial, for both duration, $F_{genotype\ x\ diet\ x\ trial}(4, 178) = 0.65$, $p = 0.63$, as well as frequency, $F_{genotype\ x\ diet\ x\ trial}(4, 178) = 0.16$, $p = 0.96$.

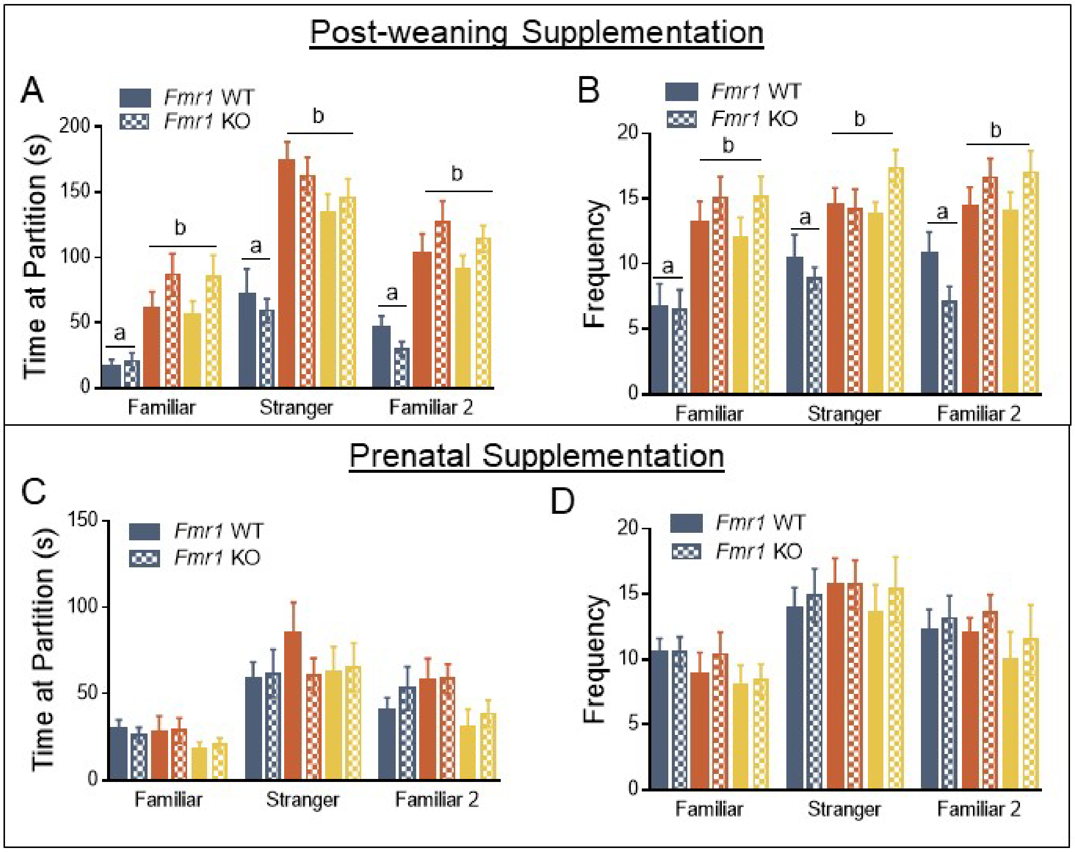


*Figure S5*. Exposure to a high fat diet during the post-weaning paradigm significantly increases frequency and duration of visits to the partition across all three trials, while the prenatal exposure has no effect on sociability. A. Post-weaning exposure to both omega-3 and control fat diets significantly increased duration of visits to the partition. B. Similarly, post-weaning exposure to both experimental diets increased the frequency of visits to the partition. C. However, prenatal exposure had no impact on duration of visits to the partition. D. The frequency of visits was also unaffected by both genotype and dietary manipulations. Data are expressed as mean ± SEM. A designation of “b” indicates that this group differed from the “a” comparison group at the level of $p < 0.05$.
